# Predicting neutralization susceptibility to combination HIV-1 monoclonal broadly neutralizing antibody regimens

**DOI:** 10.1101/2023.12.14.571616

**Authors:** Brian D. Williamson, Liana Wu, Yunda Huang, Aaron Hudson, Peter B. Gilbert

## Abstract

Combination monoclonal broadly neutralizing antibodies (bnAbs) are currently being developed for preventing HIV-1 infection. Recent work has focused on predicting in vitro neutralization potency of both individual bnAbs and combination regimens against HIV-1 pseudoviruses using Env sequence features. To predict in vitro combination regimen neutralization potency against a given HIV-1 pseudovirus, previous approaches have applied mathematical models to combine individual-bnAb neutralization and have predicted this combined neutralization value; we call this the combine-then-predict (CP) approach. However, prediction performance for some individual bnAbs has exceeded that for the combination, leading to another possibility: combining the individual-bnAb predicted values and using these to predict combination regimen neutralization; we call this the predict-then-combine (PC) approach. We explore both approaches in both simulated data and data from the Los Alamos National Laboratory’s Compile, Neutralize, and Tally NAb Panels repository. The CP approach is superior to the PC approach when the neutralization outcome of interest is binary (e.g., neutralization susceptibility, defined as inhibitory concentration *<* 1 µg/mL. For continuous outcomes, the CP approach performs at least as well as the PC approach, and is superior to the PC approach when the individual-bnAb prediction algorithms have poor performance. This knowledge may be used when building prediction models for novel antibody combinations in the absence of in vitro neutralization data for the antibody combination; this, in turn, will aid in the evaluation and down-selection of these antibody combinations into prevention efficacy trials.

## 1 Author Summary

The Antibody Mediated Prevention trials provided evidence that an antibody that can neutralize most HIV-1 viruses (a “broadly neutralizing antibody”) can prevent infection by susceptible HIV-1 viruses. Interventions containing multiple broadly neutralizing antibodies may be more effective at preventing HIV-1 infection. Understanding how well different “combination regimens” neutralize circulating HIV-1 viruses is important for designing these new interventions. However, neutralization data can be difficult to obtain; often, it is easier to obtain information about the outer envelope (Env) protein of HIV-1. Several models have been built which have good performance for predicting HIV-1 neutralization against single antibodies, but combination regimens are more difficult to study because there is limited historical neutralization data for these combinations. We investigated two approaches to using individual-antibody neutralization data to build models predicting combination neutralization: combine-then-predict and predict-then-combine. The first approach combines the individual-antibody neutralization values using validated mathematical models, and trains a model to predict this value. The second approach trains models for each individual antibody and uses the mathematical models to combine the predictions. We found in simulated experiments and using publicly-available data that the combine-then-predict approach performs at least as well as the predict-then-combine approach, and is superior in some settings. This knowledge may be used when building prediction models for novel antibody combinations in the absence of in vitro neutralization data for the antibody combination; this, in turn, will aid in the evaluation and down-selection of these antibody combinations into prevention efficacy trials.

## 2 Introduction

Monoclonal broadly neutralizing antibody (bnAb) regimens for HIV-1 prevention have been researched extensively (Mahomed et al., 2021; Julg and Barouch, 2021; Karuna and Corey, 2020; Walsh and Seaman, 2021). The Antibody Mediated Prevention (AMP) randomized efficacy trials of VRC01 versus placebo (HVTN 704/HPTN 085 and HVTN 703/HPTN 081, NCT02716675 and NCT02568215, respectively) provided evidence that a bnAb can prevent HIV-1 acquisition (Corey et al., 2021). Defining a VRC01-susceptible strain as having 80% inhibitory concentration (IC_80_) *<* 1 µg/mL, the AMP trials showed that estimated prevention efficacy of VRC01 versus placebo for VRC01-susceptible strains was 75.4% (Corey et al., 2021). Estimated prevention efficacy for strains with 50% inhibitory concentration (IC_50_) *<* 1 µg/mL was approximately 50% (Corey et al., 2021).

Several methods have been developed for predicting individual (Hepler et al., 2014; Buiu et al., 2016; Hake and Pfeifer, 2017; Rawi et al., 2019; Conti and Karplus, 2019; Yu et al., 2019; Magaret et al., 2019; Dănăilă and Buiu, 2022) and both individual and combination (Williamson et al., 2021) in vitro neutralization outcomes from Env sequence data. Many of these algorithms were trained using the Los Alamos National Laboratory’s Compile, Analyze, and Tally NAb Panels (CATNAP) database (Yoon et al., 2015) and yield good prediction performance for individual bnAbs. One limitation of CATNAP is that while sufficient numbers of pseudoviruses have measured neutralization against individual bnAbs, far fewer (and in many cases, zero) have measured neutralization against combination regimens. Additionally, many more pseudoviruses tend to have measurements of IC_50_ than IC_80_. To maximize the amount of information available when investigating combination regimens, combination neutralization is often defined as a function of the individual neutralization values, and then used as an outcome for training the prediction model (Wagh et al., 2016, 2018; Williamson et al., 2021). Other outcomes, including *multiple susceptibility* – defined as the binary indcator that *k* bnAbs in the regimen have IC_80_ *<* 1 µg/mL – could be used instead. These prediction models may be used to compare bnAb regimens when determining which to pursue in further research (Dănăilă and Buiu, 2022; Williamson et al., 2023), since neutralization is associated with prevention efficacy (Corey et al., 2021).

However, prediction performance for some individual bnAbs has exceeded that for the combination (Williamson et al., 2023). This suggests a second possible method for predicting combination neutralization outcomes: combining the individual-bnAb predicted values and using these combinations to predict neutralization by the combination regimen. In this article, we explore this approach and contrast it with the previous approach of directly predicting the combined neutralization. We study the performance of both approaches in simulated data and data from CATNAP.

## 3 Methods

### 3.1 Combining individual-bnAb neutralization values

The first method of predicting combination bnAb regimen susceptibility that we consider, which we refer to as pre-prediction combination or the *combine-then-predict* (CP) approach, involves applying a mathematical model to combine measured in vitro inhibitory concentration prior to training a prediction model. We consider two mathematical models defined by Wagh et al. (2016) for combining the in vitro neutralization values from *J* constituent bnAbs in a bnAb combination regimen: an additive model,

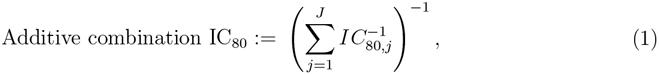

and a Bliss-Hill model,

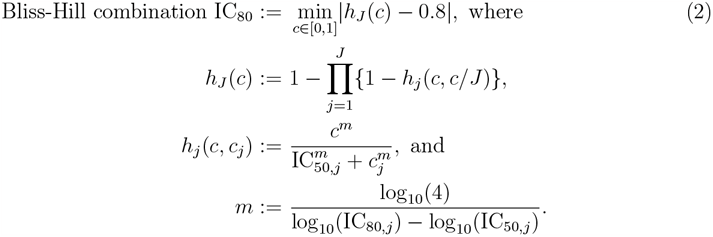

The Bliss-Hill solution is obtained using the method of Brent (1971). Combination susceptibility based on IC_80_, a binary outcome, is defined as the binary indicator that combination IC_80_ *<* 1 µg/mL. Combination IC_50_ (the 50% inhibitory concentration) and combination susceptibility based on IC_50_ are defined in a similar manner. Then, a prediction model can be trained by using the Env features to predict the combination neutralization outcome (either continuous or binary). This approach is laid out in the left-hand column of Figure 1.

**Figure 1.**
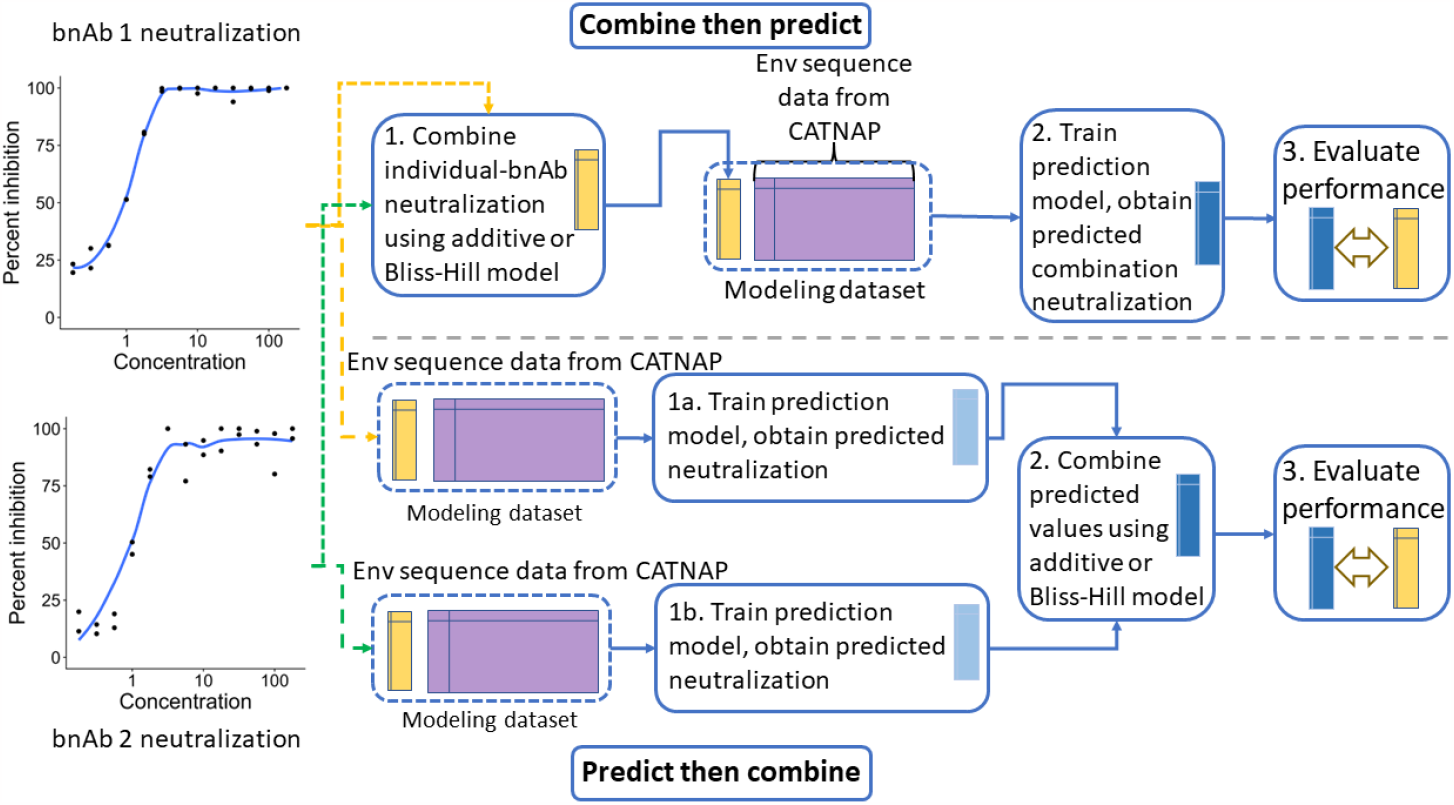
An illustration of the combine-then-predict (top) and predict-then-combine (bottom) approaches to predicting combination regimen in vitro neutralization, for a two-bnAb combination regimen.

One could instead perform post-prediction combination, which is the second method of predicting combination bnAb regimen susceptibility that we consider. We will refer to this as the *predict-then-combine* (PC) approach. This method involves first training *J* prediction models using the Env features to predict the continuous neutralization outcome (e.g., IC_80_) of each individual bnAb in the combination regimen. We then obtain predicted neutralization 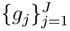 and use either the additive or Bliss-Hill model to combine the predicted values, using the predictions *g*_*j*_ in place of the measured IC_80,*j*_ or IC_50,*j*_. To predict susceptibility, we check if the combined predicted value is less than 1 µg/mL.

### 3.2 CATNAP datasets

We consider several bnAb combination regimens with data available in CATNAP. These regimens consist of those undergoing HIV Vaccine Trials Network (HVTN) or HIV Prevention Trials Network (HPTN) clinical testing as of October 2022 (Williamson et al., 2023) and those with at least 125 pseudoviruses with direct measurement of neutralization by the bnAb combination (Williamson et al., 2021). The full list of bnAb combination regimens is provided in Table 1.

**Table 1:** Broadly neutralizing antibody combination regimens either with at least 125 sequences in CATNAP with direct measurement of combination neutralization or undergoing HVTN/HPTN clinical testing as of October 2022.

| Combination regimen | Number of sequences in CATNAP with $IC_{80}$ data measured against each constituent bnAb | Number of sequences in CATNAP with $IC_{80}$ data measured against the combination |
| --- | --- | --- |
| BG1 + BG18 + NC37 | 118 | 119 |
| PG9 + PGT128 | 300 | 125 |
| PG9 + PGT128 + VRC07 | 300 | 125 |
| PG9 + VRC07 | 392 | 125 |
| PGT128 + VRC07 | 299 | 125 |
| VRC07-523-LS + 10-1074 | 400 | 0 |
| VRC07-523-LS + PGDM1400 | 400 | 0 |
| VRC07-523-LS + PGT121 | 400 | 0 |
| VRC07-523-LS + PGDM1400 + PGT121 | 400 | 0 |
| VRC07-523-LS + VRC26.25 | 400 | 0 |
| 3BNC117 + PG9 | 400 | 125 |
| 10-1074 + 3BNC117 | 542 | 125 |
| 10-1074 + 3BNC117 + PG9 | 402 | 125 |
| 10-1074 + 10E8 | 403 | 125 |
| 10-1074 + 10E8 + 3BNC117 | 403 | 125 |
| 10-1074 + PG9 | 402 | 125 |
| 10E8 + 3BNC117 | 400 | 125 |
| 10E8 + 3BNC117 + PG9 | 400 | 125 |
| 10E8 + PG9 + PGT128 | 300 | 125 |
| 10E8 + PG9 + VRC07 | 392 | 125 |
| 10E8 + PGT128 + VRC07 | 300 | 125 |

We began by using the Super LeArner Prediction of NAb Panels (SLAPNAP) tool (Williamson et al., 2021) to download and format a dataset for each unique bnAb in Table 1 consisting of individual-bnAb IC_50_ and IC_80_ values, along with Env gp160 sequence features. We also downloaded and formatted datasets of directly measured combination neutralization for those bnAb regimens in Table 1 with such information (Table 1 column 3 not equal to zero). The features for each pseudovirus consist of geographic information (binary indicator variables describing the geographic region of origin), viral geometry variables (length, number of sequons, and number of cysteines in the Env, gp120, V2, V3, and V5 regions), and amino acid sequence variables (binary indicators of residues containing amino acids, frameshifts, gaps, stops, or sequons at each HXB2-referenced site in gp160). Each dataset had approximately 6000 columns.

We next merged the individual-bnAb neutralization values and directly-observed combination neutralization (if available) into a single dataset for each bnAb regimen in Table 1. We computed model-predicted combination neutralization using both the additive and Bliss-Hill models described above. The statistical learning approaches described below aimed to predict the following neutralization outcomes for each individual bnAb in the regimen: (1) the continuous log_10_ IC_50_; (2) the continuous log_10_ IC_80_; (3) the dichotomous outcome IC_50_ susceptibility, defined as IC_50_ *<* 1 µg/mL; and (4) the dichotomous outcome IC_80_ susceptibility, defined as IC_80_ *<* 1 µg/mL. We also aimed to predict the following outcomes for each combination regimen and both the additive and Bliss-Hill combination models: (5) the continuous log_10_ combination IC_50_; (6) the continuous log_10_ combination IC_80_; (7) the dichotomous outcome combination IC_50_ susceptibility, defined as combination IC_50_ *<* 1 µg/mL; and (8) the dichotomous outcome combination IC_80_ susceptibility, defined as combination IC_80_ *<* 1 µg/mL. For combination regimens with directly-observed combination neutralization, we further aimed to predict outcomes (1) through (4) but using the directly-observed combination neutralization values.

## 4 Results

### 4.1 Numerical experiments

In this section, we performed a simple numerical experiment to illustrate the prediction performance of the CP versus PC approach for predicting the combination neutralization value or susceptibility. For sample size *n ∈ {*100, 200, …, 1000*}* and three bnAbs, we generated 1000 Env AA sequence features and IC_80_ values for each bnAb and each of the *n* simulated HIV-1 pseudoviruses. In our simulated setting, only ten Env AA sequence features impact the IC_80_. We considered two scenarios: one where there was a weak relationship between these ten Env AA sequence features and IC_80_, and one where there was a strong relationship. The Env AA sequence features *X* were generated by first creating a latent matrix *W* using a multivariate normal distribution with identity covariance matrix; the first six Env features are equal to the corresponding latent value, while the remaining are binary, with *X*_*ik*_ = *I{W*_*ik*_ *< q*_40_(*W*_*k*_)*}* for *i* = 1, …, *n, k ∈ {*6, …, 1000*}* and *q*_*δ*_(*v*) denoting the *δ*-quantile of *v*. The IC_80_ values were generated according to IC_80,*j*_ = *Xβ*_*j*_, where *j ∈ {*1, 2, 3*}* denotes the bnAb and *β*_*j*_ = [*α*_*j*_**1**_10_, **0**_*p−*10_]^*T*^. In the weak-relationship scenario, *α*_1_ = 0.2, *α*_2_ = 0.4, and *α*_3_ = 0.1, resulting in proportion of

HIV-1 pseudoviruses susceptible to each of the three bnAbs equal to 0.36, 0.29, and 0.42, respectively. In the strong-relationship scenario, *α*_1_ = 1, *α*_2_ = 2, and *α*_3_ = 0.5, resulting in proportion equal to 0.23, 0.21, and 0.27, respectively. We then obtained combined IC_80_ using the additive model of Wagh et al. (2016) (see Methods). The proportion susceptible using combination IC_80_ was 0.74 in the weak-relationship scenario and 0.38 in the strong-relationship scenario. We then used the lasso (Tibshirani, 1996) to (i) predict the individual-bnAb neutralization (both continuous and binary); (ii) predict the combination neutralization directly (both continuous and binary), i.e., use the CP approach; and (iii) combine the continuous-valued predictions from (i) using the additive model, i.e., use the PC approach. We used 10-fold cross-validation to obtain the optimal lasso tuning parameter in all cases.

We assessed prediction performance using an additional layer of 10-fold cross-validation, where the lasso was trained on nine-tenths of the data and prediction performance was estimated on the remaining tenth. We evaluated prediction performance using cross-validated (CV) R-squared for continuous outcomes and cross-validated area under the receiver operating characteristic curve (CV AUC) for binary outcomes. We also obtained a 95% confidence interval for the prediction performance (Williamson et al., 2021; LeDell et al., 2015). We repeated the above data-generating, prediction, and prediction-performance process 2500 times for each sample size, and computed the average point estimate, empirical standard error of the point estimates, and average confidence interval width.

We display the results for these simulated data in Figures 2 and 3. In Figure 2 (left-hand column), we see that the prediction performance varies across individual bnAbs in the combination regimen, with the second bnAb having uniformly highest prediction performance; the third bnAb having uniformly lowest prediction performance; and the first bnAb having prediction performance between the other two. Prediction performance is degraded in the weak-relationship case, which is expected. Confidence interval width decreases with increasing sample size for both outcomes, as expected (right-hand column of Figure 2), though width tends to be higher in the weak-relationship case than in the strong-relationship case. In Figure 3 (left-hand column), we see that prediction performance is similar across the two methods of predicting combination neutralization for the continuous outcome (nearly within Monte-Carlo error) in the strong-relationship case, but is uniformly higher when using the CP approach in the weak-relationship case, corresponding to a case where CV R-squared for all individual bnAbs was less than 0.5. Training prediction models using the CP approach results in higher prediction performance for susceptibility regardless of the strength of relationship between predictors and outcome. Confidence interval width is uniformly smaller for the CP method (Figure 3 right-hand column).

**Figure 2.**
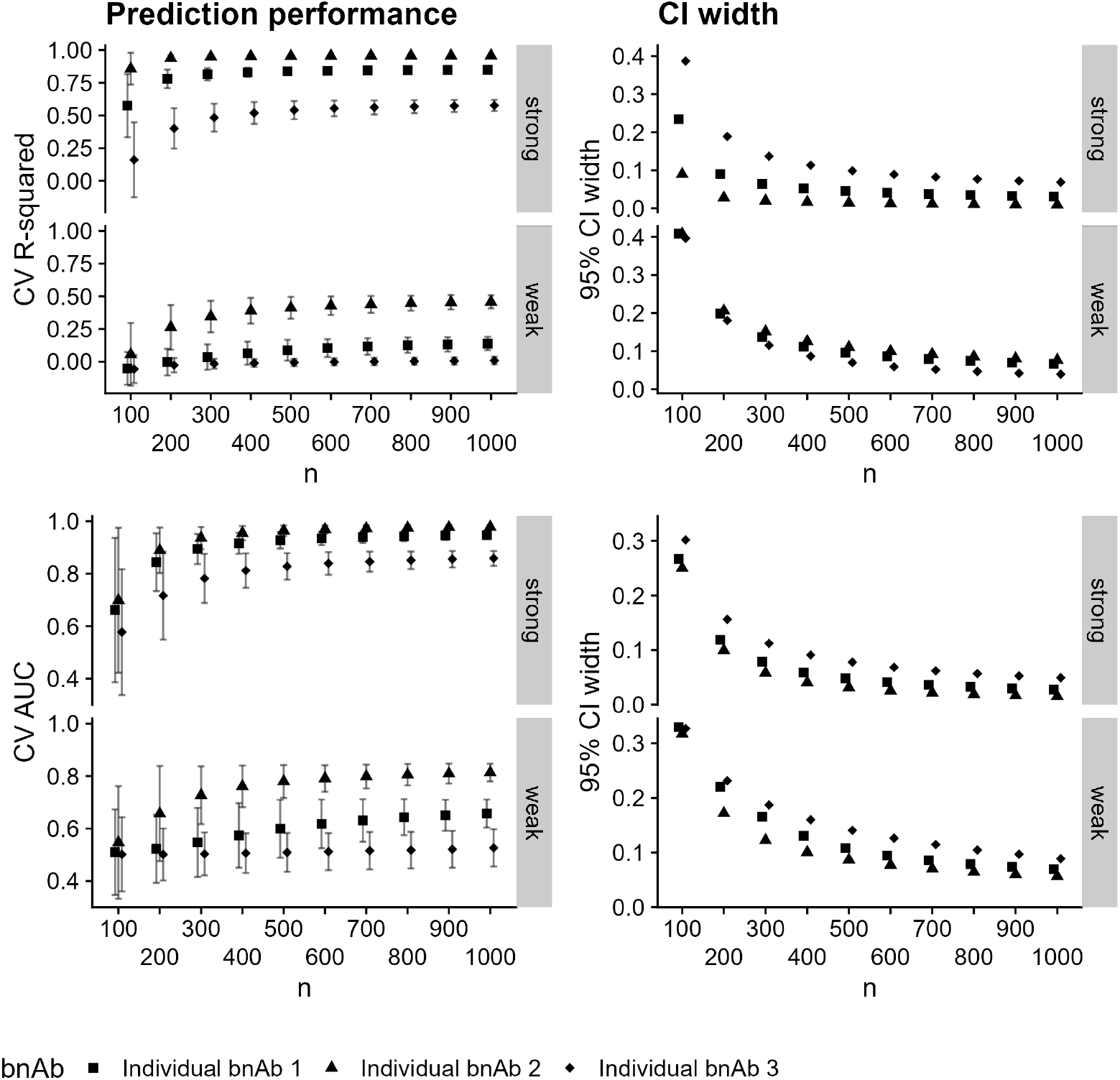
Prediction performance (left-hand column) and 95% confidence interval width (right-hand column) versus sample size for predicting IC_80_ (top row) and IC_80_ *<* 1 µg/mL for each of three individual bnAbs averaged over 2500 simulated datasets. Panels within rows denote a strong or weak relationship between the Env AA predictors and the outcome. Shapes denote the individual bnAbs, and Monte-Carlo error is displayed in error bars.

**Figure 3.**
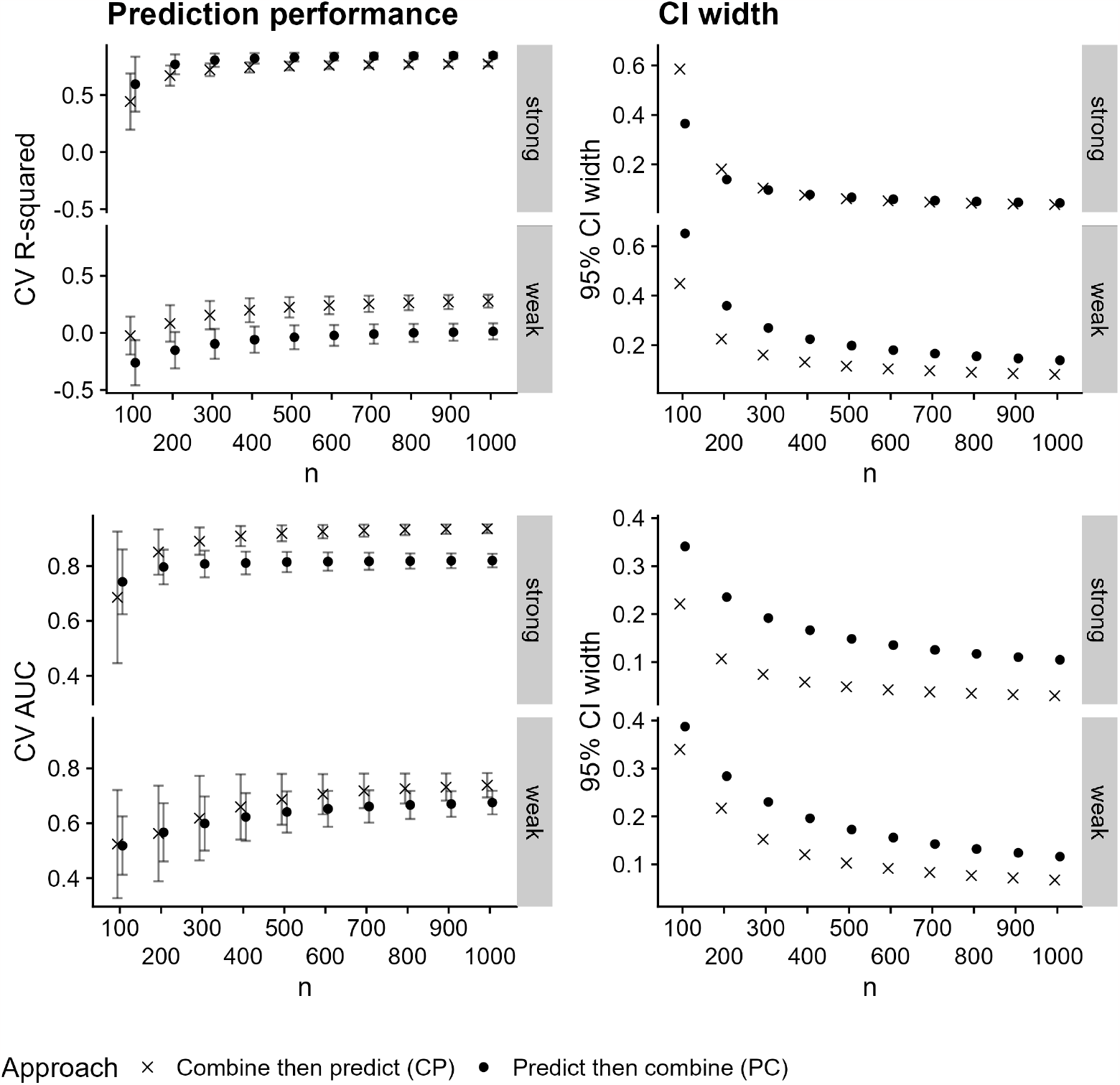
Prediction performance (left-hand column) and 95% confidence interval width (right-hand column) versus sample size for predicting combination IC_80_ (top row) and combination IC_80_ *<* 1 µg/mL using both the CP and PC approaches, averaged over 2500 simulated datasets. Panels within rows denote a strong or weak relationship between the Env AA predictors and the outcome. Shapes denote the approach, and Monte-Carlo error is displayed in error bars.

### 4.2 Results from bnAb combinations in CATNAP

We further explored the distinction between the CP and PC approaches by obtaining prediction performance for the bnAb combination regimens in Table 1. We followed the approach described in Methods to define the dataset for each bnAb regimen.

For each bnAb regimen in Table 1, we followed the same procedure as in the previous section: we estimated prediction performance for both continuous and binary outcomes (both based on IC_50_ and IC_80_) using ten-fold cross-validation, with additional ten-fold cross-validation to select the lasso tuning parameters. As before, we evaluated both individual-bnAb prediction performance and performance for predicting combination regimen neutralization by either combining the individual-bnAb predictions or by directly predicting the combination neutralization. In this case, we considered both the additive and Bliss-Hill models (Wagh et al., 2016, see Methods) for combining neutralization values.

For the bnAb regimens with direct lab-measured neutralization against the combination regimen (third column of Table 1), we further assessed the performance of both approaches for predicting these neutralization values. Below, we highlight the results for two bnAb regimens, VRC07-523-LS + 10-1074 (chosen because it is a regimen in clinical testing) and 10-1074 + 10E8 (chosen because it is a regimen with direct measurement of neutralization against the regimen). Since predicting the directly-measured neutralization against a combination regimen is an important goal, in cases where this direct measurement exists we evaluate prediction performance for this outcome as well as the model-combined outcome.

The proportion of pseudoviruses susceptible (measured using IC_80_) to VRC07-523-LS, 10-1074, and 10E8 are 78.75%, 48.89%, and 21.34%, respectively. The proportion estimated to be sensitive to VRC07-523-LS + 10-1074 is 89.5% and 93.5% for the additive and Bliss-Hill models, respectively. The proportion estimated to be sensitive to 10-1074 + 10E8 is 59.65% and 79.7% for the additive and Bliss-Hill models, respectively, while the true proportion susceptible is 64.8%. Based on the number of pseudoviruses, the number of susceptible viruses is comparable to *n ∈ {*600, …, 1000*}* in the simulations above. Results for the remaining bnAb regimens from Table 1 can be found in Figures S1–S18.

We display the results for predicting neutralization susceptibility to VRC07-523-LS + 10-1074 in Figure 4. In the left-hand column, we see that prediction performance is higher for 10-1074 for both continuous and binary neutralization outcomes. In the right-hand column, we see that for both continuous and binary outcomes, the CP approach leads to better prediction performance. This is particularly striking for the Bliss-Hill model in the continuous-outcome setting. In this case, we do not have direct measurements of neutralization by the combination regimen. As in the simulations, prediction performance for the combination tends to be less than the best individual-bnAb prediction performance. When using the CP approach, prediction performance for the combination tends to be greater than the worst individual-bnAb prediction performance.

**Figure 4.**
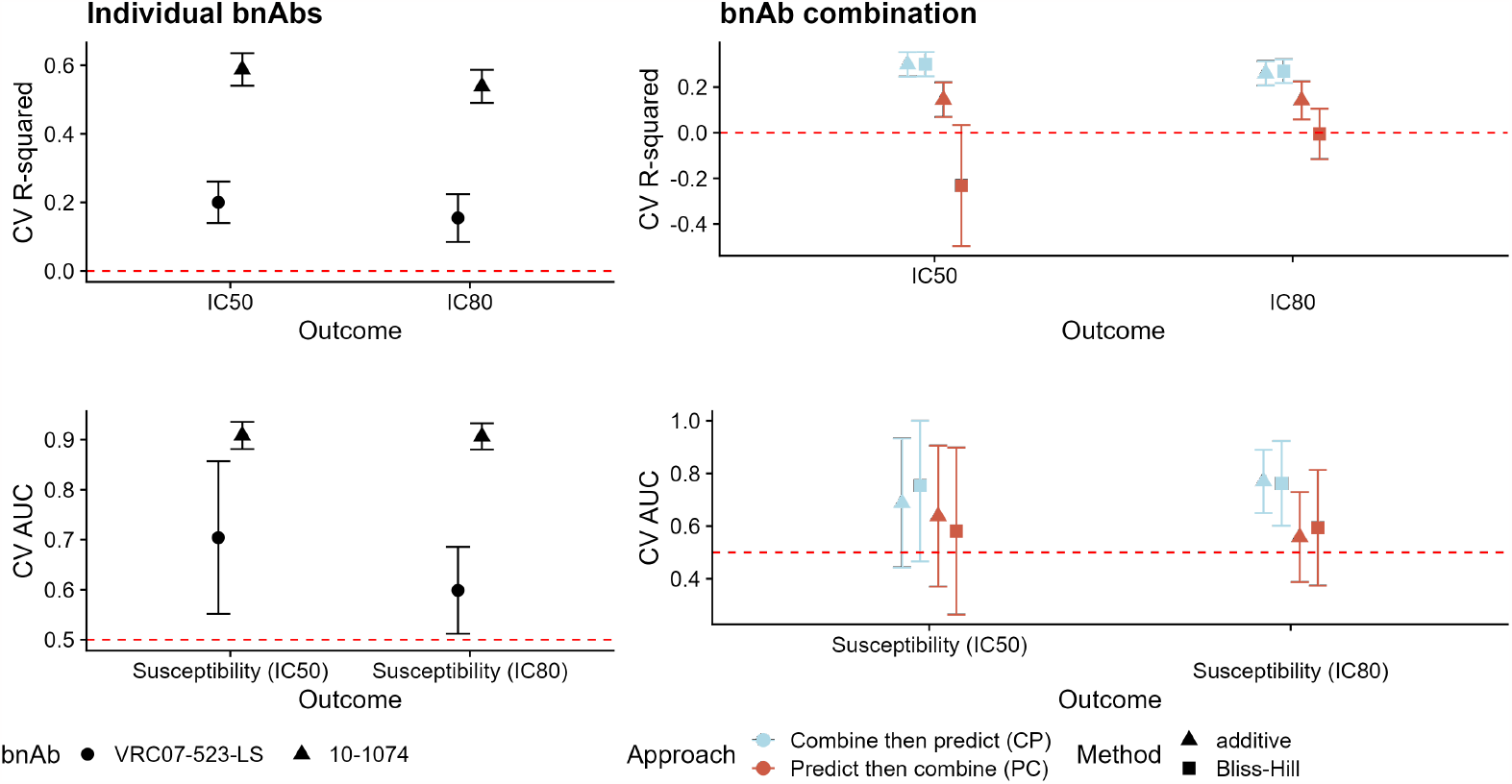
Prediction performance for continuous (top row, CV R-squared) and binary (bottom row, CV AUC) neutralization outcomes for individual bnAbs (left-hand column) and the combination (right-hand column) VRC07-523-LS + 10-1074. For individual bnAbs, prediction performance is evaluated against the observed IC_50_ or IC_80_ values for the given bnAb; shapes denote the bnAb. For combination bnAbs, prediction performance is evaluated against the calculated combination IC_50_ or IC_80_ values based on the observed bnAb-specific values using the additive or Bliss-Hill method; shapes denote the combination method (additive or Bliss-Hill) and color denotes the approach (CP or PC). Error bars reflect 95% confidence intervals.

**Figure 5.**
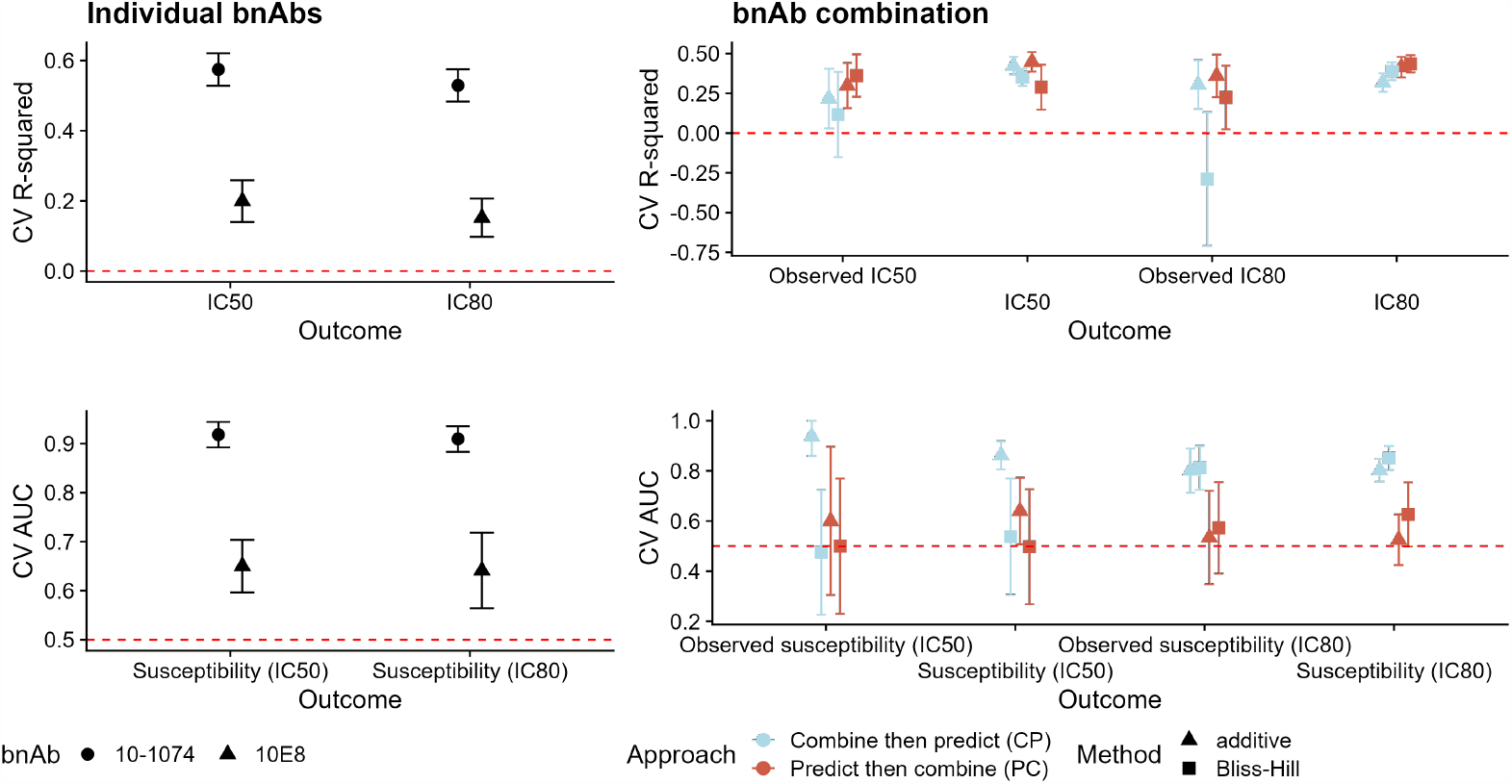
Prediction performance for continuous (top row, CV R-squared) and binary (bottom row, CV AUC) neutralization outcomes for individual bnAbs (left-hand column) and the combination (right-hand column) 10-1074 + 10E8. For individual bnAbs, prediction performance is evaluated against the observed IC_50_ or IC_80_ values for the given bnAb; shapes denote the bnAb. For combination bnAbs, prediction performance is evaluated against both the observed IC_50_ or IC_80_ values based on the bnAb regimen (denoted by the prefix “observed”) and the calculated combination IC_50_ or IC_80_ values based on the observed bnAb-specific values using the additive or Bliss-Hill method; shapes denote the combination method (additive or Bliss-Hill) and color denotes the approach (CP or PC). Error bars reflect 95% confidence intervals.

We display the results for predicting neutralization susceptibility to 10-1074 + 10E8 in Figure 4. In the left-hand column, we see that prediction performance is again higher for 10-1074 for both continuous and binary neutralization outcomes. In the right-hand column, we see that prediction performance for continuous outcomes is similar between the CP and PC approaches; for binary outcomes, performance is better when using the CP approach. In this case, we also have direct measurements of neutralization by the combination regimen, and the results follow a similar pattern to the combination neutralization results. These results are similar to the results from the simulations.

Results for the remaining bnAb regimens from Table 1 followed similar patterns (Figures S1– S18). We observed that prediction performance for binary neutralization outcomes (e.g., IC_80_ *<* 1 µg/mL) was better when using the CP approach than when using the PC approach. Results for continuous outcomes are mixed. However, the results reflect what we saw in the simulations: when continuous-outcome prediction performance for the individual bnAbs is higher than 0.5 (e.g., for 10-1074 + 10E8 above, or 10-1074 + PG9, Figure S14) then prediction performance is similar using either the CP or PC approach. For many combinations, however, the CP approach results in better prediction performance than the PC approach.

## 5 Discussion

When predicting neutralization susceptibility to a combination bnAb regimen, a key consideration is whether to directly predict the combination neutralization susceptibility (combined using the individual-bnAb neutralization values), which we called the combine-then-predict (CP) approach, or to combine the predictions of individual-bnAb neutralization, which we called the predict-then-combine (PC) approach. In both simulated experiments and our data analysis, we found that prediction performance for binary neutralization outcomes (e.g., IC_80_ *<* 1 µg/mL) was better using the CP approach compared to the PC approach. Results for continuous outcomes were mixed: in the simulations and the data analysis, we observed that in small samples, prediction performance for combination neutralization was similar between the two approaches. In larger samples (in the simulations), the PC approach seems to result in slightly better prediction performance, when individual-model prediction performance is high. This knowledge may be used when building prediction models for novel antibody combinations in the absence of in vitro neutralization data for the antibody combination; this, in turn, will aid in the evaluation and down-selection of these antibody combinations into prevention efficacy trials.

It is intuitive that increased individual-model prediction performance leads to increased PC performance, since as we noted above prediction errors may be compounded in this approach. The CP approach is similar to averaging IC_80_ values over technical replicates, which generally improves the signal-to-noise ratio (SNR). This is similar to the phenomenon observed in Huang et al. (2017), where the authors observed that averaging several immune response endpoints prior to ranking vaccine regimens resulted in better selection of vaccine regimens than ranking the immune responses and then averaging the ranks. The method of Follmann (2018) has also been used to compare multiple immune responses (see, e.g., Benkeser et al., 2023), and considers the SNR for each immune response. Applying this method to the simulated examples yields that the SNR for the combination susceptibility is always higher than the SNR for the individual-bnAb susceptibilities, while the SNR for the combination IC_80_ is only higher than the SNR for the individual-bnAb IC_80_ values in the case where the individual-bnAb prediction performance is high; the PC approach performed better than the CP approach in this case. In cases with high SNR for the individual bnAb immune responses, denoising through averaging (i.e., using the CP approach) may be less important. This suggests that the SNR can be used to reliably predict whether the CP or PC approach may perform better in practice. However, in all cases, the CP approach led to narrower confidence intervals for prediction performance.

These results appear to be more pronounced for the Bliss-Hill model than for the additive model. This may be due in part to the fact that the Bliss-Hill model uses both IC_50_ and IC_80_, so any prediction error in the individual-bnAb models is compounded; the additive model relies only on predictions of either IC_50_ or IC_80_.

This study has several limitations. The first is that we only used a single prediction method, the lasso, for our analysis of the CATNAP data. Several groups of authors have observed that either an ensemble of multiple prediction methods (Williamson et al., 2021) or more flexible prediction methods (Hake and Pfeifer, 2017; Rawi et al., 2019; Dănăilă and Buiu, 2022) can improve prediction performance using data from CATNAP. While we could certainly observe increased prediction performance by using a more flexible prediction method, comparisons of the CP and PC approaches within a bnAb regimen are nonetheless valid since the predicted values from each approach are evaluated in the same manner regardless of the prediction model used to obtain these values. Additionally, as we saw in the simulations, a CV R-squared greater than approximately 0.5 for all bnAbs in a regimen is necessary for the PC approach to outperform the CP approach; in previous work using an ensemble prediction method, we observed very few CV R-squared values greater than 0.5 across a large number of individual bnAbs (Williamson et al., 2021). Second, all of the predictions in this work are based only on the HIV-1 Env sequence and do not account for the bnAb variable region amino acid sequence, which has been observed to increase prediction performance (Dănăilă and Buiu, 2022). Finally, in the simulations, the lasso model provides an unbiased estimator of the true data-generating model, so it is unlikely that prediction performance could be improved. A second limitation is that there is relatively little neutralization data in CATNAP on combination regimens. This limits our ability to either develop prediction models using the neutralization outcomes based on the combination regimen (here, we held these viruses out as an independent test set) or to obtain precise estimates of prediction performance.

## Acknowledgements

This work was supported by the National Institute of Allergy and Infectious Diseases (NIAID) of the National Institutes of Health (NIH) U.S. Public Health Service Grant UM1AI068635 and by the National Library of Medicine (NLM) of the NIH grant R25LM014210. The content is solely the responsibility of the authors and does not necessarily represent the official views of the NIH.

## Supporting information

Figure S1. Prediction performance for continuous (top row, CV R-squared) and binary (bottom row, CV AUC) neutralization outcomes for individual bnAbs (left-hand column) and the combination (right-hand column) BG1 + BG18 + NC37. For individual bnAbs, prediction performance is evaluated against the observed IC_50_ or IC_80_ values for the given bnAb; shapes denote the bnAb. For combination bnAbs, prediction performance is evaluated against both the observed IC_50_ or IC_80_ values based on the bnAb regimen (denoted by the prefix “observed”) and the calculated combination IC_50_ or IC_80_ values based on the observed bnAb-specific values using the additive or Bliss-Hill method; shapes denote the combination method (additive or Bliss-Hill) and color denotes the approach (CP or PC). Error bars reflect 95% confidence intervals.

Figure S2. Prediction performance for continuous (top row, CV R-squared) and binary (bottom row, CV AUC) neutralization outcomes for individual bnAbs (left-hand column) and the combination (right-hand column) PG9 + PGT128. For individual bnAbs, prediction performance is evaluated against the observed IC_50_ or IC_80_ values for the given bnAb; shapes denote the bnAb. For combination bnAbs, prediction performance is evaluated against both the observed IC_50_ or IC_80_ values based on the bnAb regimen (denoted by the prefix “observed”) and the calculated combination IC_50_ or IC_80_ values based on the observed bnAb-specific values using the additive or Bliss-Hill method; shapes denote the combination method (additive or Bliss-Hill) and color denotes the approach (CP or PC). Error bars reflect 95% confidence intervals.

Figure S3. Prediction performance for continuous (top row, CV R-squared) and binary (bottom row, CV AUC) neutralization outcomes for individual bnAbs (left-hand column) and the combination (right-hand column) PG9 + PGT128 + VRC07. For individual bnAbs, prediction performance is evaluated against the observed IC_50_ or IC_80_ values for the given bnAb; shapes denote the bnAb. For combination bnAbs, prediction performance is evaluated against both the observed IC_50_ or IC_80_ values based on the bnAb regimen (denoted by the prefix “observed”) and the calculated combination IC_50_ or IC_80_ values based on the observed bnAb-specific values using the additive or Bliss-Hill method; shapes denote the combination method (additive or Bliss-Hill) and color denotes the approach (CP or PC). Error bars reflect 95% confidence intervals.

Figure S4. Prediction performance for continuous (top row, CV R-squared) and binary (bottom row, CV AUC) neutralization outcomes for individual bnAbs (left-hand column) and the combination (right-hand column) PG9 + VRC07. For individual bnAbs, prediction performance is evaluated against the observed IC_50_ or IC_80_ values for the given bnAb; shapes denote the bnAb. For combination bnAbs, prediction performance is evaluated against both the observed IC_50_ or IC_80_ values based on the bnAb regimen (denoted by the prefix “observed”) and the calculated combination IC_50_ or IC_80_ values based on the observed bnAb-specific values using the additive or Bliss-Hill method; shapes denote the combination method (additive or Bliss-Hill) and color denotes the approach (CP or PC). Error bars reflect 95% confidence intervals.

Figure S5. Prediction performance for continuous (top row, CV R-squared) and binary (bottom row, CV AUC) neutralization outcomes for individual bnAbs (left-hand column) and the combination (right-hand column) PGT128 + VRC07. For individual bnAbs, prediction performance is evaluated against the observed IC_50_ or IC_80_ values for the given bnAb; shapes denote the bnAb. For combination bnAbs, prediction performance is evaluated against both the observed IC_50_ or IC_80_ values based on the bnAb regimen (denoted by the prefix “observed”) and the calculated combination IC_50_ or IC_80_ values based on the observed bnAb-specific values using the additive or Bliss-Hill method; shapes denote the combination method (additive or Bliss-Hill) and color denotes the approach (CP or PC). Error bars reflect 95% confidence intervals.

Figure S6. Prediction performance for continuous (top row, CV R-squared) and binary (bottom row, CV AUC) neutralization outcomes for individual bnAbs (left-hand column) and the combination (right-hand column) VRC07-523-LS + PGDM1400. For individual bnAbs, prediction performance is evaluated against the observed IC_50_ or IC_80_ values for the given bnAb; shapes denote the bnAb. For combination bnAbs, prediction performance is evaluated against the calculated combination IC_50_ or IC_80_ values based on the observed bnAb-specific values using the additive or Bliss-Hill method; shapes denote the combination method (additive or Bliss-Hill) and color denotes the approach (CP or PC). Error bars reflect 95% confidence intervals.

Figure S7. Prediction performance for continuous (top row, CV R-squared) and binary (bottom row, CV AUC) neutralization outcomes for individual bnAbs (left-hand column) and the combination (right-hand column) VRC07-523-LS + PGT121. For individual bnAbs, prediction performance is evaluated against the observed IC_50_ or IC_80_ values for the given bnAb; shapes denote the bnAb. For combination bnAbs, prediction performance is evaluated against the calculated combination IC_50_ or IC_80_ values based on the observed bnAb-specific values using the additive or Bliss-Hill method; shapes denote the combination method (additive or Bliss-Hill) and color denotes the approach (CP or PC). Error bars reflect 95% confidence intervals.

Figure S8. Prediction performance for continuous (top row, CV R-squared) and binary (bottom row, CV AUC) neutralization outcomes for individual bnAbs (left-hand column) and the combination (right-hand column) VRC07-523-LS + PGT121 + PGDM1400. For individual bnAbs, prediction performance is evaluated against the observed IC_50_ or IC_80_ values for the given bnAb; shapes denote the bnAb. For combination bnAbs, prediction performance is evaluated against the calculated combination IC_50_ or IC_80_ values based on the observed bnAb-specific values using the additive or Bliss-Hill method; shapes denote the combination method (additive or Bliss-Hill) and color denotes the approach (CP or PC). Error bars reflect 95% confidence intervals.

Figure S9. Prediction performance for continuous (top row, CV R-squared) and binary (bottom row, CV AUC) neutralization outcomes for individual bnAbs (left-hand column) and the combination (right-hand column) VRC07-523-LS + VRC26.25. For individual bnAbs, prediction performance is evaluated against the observed IC_50_ or IC_80_ values for the given bnAb; shapes denote the bnAb. For combination bnAbs, prediction performance is evaluated against the calculated combination IC_50_ or IC_80_ values based on the observed bnAb-specific values using the additive or Bliss-Hill method; shapes denote the combination method (additive or Bliss-Hill) and color denotes the approach (CP or PC). Error bars reflect 95% confidence intervals.

Figure S10. Prediction performance for continuous (top row, CV R-squared) and binary (bottom row, CV AUC) neutralization outcomes for individual bnAbs (left-hand column) and the combination (right-hand column) 3BNC117 + PG9. For individual bnAbs, prediction performance is evaluated against the observed IC_50_ or IC_80_ values for the given bnAb; shapes denote the bnAb. For combination bnAbs, prediction performance is evaluated against both the observed IC_50_ or IC_80_ values based on the bnAb regimen (denoted by the prefix “observed”) and the calculated combination IC_50_ or IC_80_ values based on the observed bnAb-specific values using the additive or Bliss-Hill method; shapes denote the combination method (additive or Bliss-Hill) and color denotes the approach (CP or PC). Error bars reflect 95% confidence intervals.

Figure S11. Prediction performance for continuous (top row, CV R-squared) and binary (bottom row, CV AUC) neutralization outcomes for individual bnAbs (left-hand column) and the combination (right-hand column) 10-1074 + 3BNC117. For individual bnAbs, prediction performance is evaluated against the observed IC_50_ or IC_80_ values for the given bnAb; shapes denote the bnAb. For combination bnAbs, prediction performance is evaluated against both the observed IC_50_ or IC_80_ values based on the bnAb regimen (denoted by the prefix “observed”) and the calculated combination IC_50_ or IC_80_ values based on the observed bnAb-specific values using the additive or Bliss-Hill method; shapes denote the combination method (additive or Bliss-Hill) and color denotes the approach (CP or PC). Error bars reflect 95% confidence intervals.

Figure S12. Prediction performance for continuous (top row, CV R-squared) and binary (bottom row, CV AUC) neutralization outcomes for individual bnAbs (left-hand column) and the combination (right-hand column) 10-1074 + 3BNC117 + PG9. For individual bnAbs, prediction performance is evaluated against the observed IC_50_ or IC_80_ values for the given bnAb; shapes denote the bnAb. For combination bnAbs, prediction performance is evaluated against both the observed IC_50_ or IC_80_ values based on the bnAb regimen (denoted by the prefix “observed”) and the calculated combination IC_50_ or IC_80_ values based on the observed bnAb-specific values using the additive or Bliss-Hill method; shapes denote the combination method (additive or Bliss-Hill) and color denotes the approach (CP or PC). Error bars reflect 95% confidence intervals.

Figure S13. Prediction performance for continuous (top row, CV R-squared) and binary (bottom row, CV AUC) neutralization outcomes for individual bnAbs (left-hand column) and the combination (right-hand column) 10-1074 + 10E8 + 3BNC117. For individual bnAbs, prediction performance is evaluated against the observed IC_50_ or IC_80_ values for the given bnAb; shapes denote the bnAb. For combination bnAbs, prediction performance is evaluated against both the observed IC_50_ or IC_80_ values based on the bnAb regimen (denoted by the prefix “observed”) and the calculated combination IC_50_ or IC_80_ values based on the observed bnAb-specific values using the additive or Bliss-Hill method; shapes denote the combination method (additive or Bliss-Hill) and color denotes the approach (CP or PC). Error bars reflect 95% confidence intervals.

Figure S14. Prediction performance for continuous (top row, CV R-squared) and binary (bottom row, CV AUC) neutralization outcomes for individual bnAbs (left-hand column) and the combination (right-hand column) 10-1074 + PG9. For individual bnAbs, prediction performance is evaluated against the observed IC_50_ or IC_80_ values for the given bnAb; shapes denote the bnAb. For combination bnAbs, prediction performance is evaluated against both the observed IC_50_ or IC_80_ values based on the bnAb regimen (denoted by the prefix “observed”) and the calculated combination IC_50_ or IC_80_ values based on the observed bnAb-specific values using the additive or Bliss-Hill method; shapes denote the combination method (additive or Bliss-Hill) and color denotes the approach (CP or PC). Error bars reflect 95% confidence intervals.

Figure S15. Prediction performance for continuous (top row, CV R-squared) and binary (bottom row, CV AUC) neutralization outcomes for individual bnAbs (left-hand column) and the combination (right-hand column) 10E8 + 3BNC117. For individual bnAbs, prediction performance is evaluated against the observed IC_50_ or IC_80_ values for the given bnAb; shapes denote the bnAb. For combination bnAbs, prediction performance is evaluated against both the observed IC_50_ or IC_80_ values based on the bnAb regimen (denoted by the prefix “observed”) and the calculated combination IC_50_ or IC_80_ values based on the observed bnAb-specific values using the additive or Bliss-Hill method; shapes denote the combination method (additive or Bliss-Hill) and color denotes the approach (CP or PC). Error bars reflect 95% confidence intervals.

Figure S16. Prediction performance for continuous (top row, CV R-squared) and binary (bottom row, CV AUC) neutralization outcomes for individual bnAbs (left-hand column) and the combination (right-hand column) 10E8 + 3BNC117 + PG9. For individual bnAbs, prediction performance is evaluated against the observed IC_50_ or IC_80_ values for the given bnAb; shapes denote the bnAb. For combination bnAbs, prediction performance is evaluated against both the observed IC_50_ or IC_80_ values based on the bnAb regimen (denoted by the prefix “observed”) and the calculated combination IC_50_ or IC_80_ values based on the observed bnAb-specific values using the additive or Bliss-Hill method; shapes denote the combination method (additive or Bliss-Hill) and color denotes the approach (CP or PC). Error bars reflect 95% confidence intervals.

Figure S17. Prediction performance for continuous (top row, CV R-squared) and binary (bottom row, CV AUC) neutralization outcomes for individual bnAbs (left-hand column) and the combination (right-hand column) 10E8 + PG9 + PGT128. For individual bnAbs, prediction performance is evaluated against the observed IC_50_ or IC_80_ values for the given bnAb; shapes denote the bnAb. For combination bnAbs, prediction performance is evaluated against both the observed IC_50_ or IC_80_ values based on the bnAb regimen (denoted by the prefix “observed”) and the calculated combination IC_50_ or IC_80_ values based on the observed bnAb-specific values using the additive or Bliss-Hill method; shapes denote the combination method (additive or Bliss-Hill) and color denotes the approach (CP or PC). Error bars reflect 95% confidence intervals.

Figure S18. Prediction performance for continuous (top row, CV R-squared) and binary (bottom row, CV AUC) neutralization outcomes for individual bnAbs (left-hand column) and the combination (right-hand column) 10E8 + PGT128 + VRC07. For individual bnAbs, prediction performance is evaluated against the observed IC_50_ or IC_80_ values for the given bnAb; shapes denote the bnAb. For combination bnAbs, prediction performance is evaluated against the calculated combination IC_50_ or IC_80_ values based on the observed bnAb-specific values using the additive or Bliss-Hill method; shapes denote the combination method (additive or Bliss-Hill) and color denotes the approach (CP or PC). Error bars reflect 95% confidence intervals.

## References

Benkeser, D., D. C. Montefiori, A. B. McDermott, Y. Fong, H. E. Janes, W. Deng, H. Zhou, C. R. Houchens, K. Martins, L. Jayashankar, et al. (2023). Comparing antibody assays as correlates of protection against COVID-19 in the COVE mRNA-1273 vaccine efficacy trial. Science Translational Medicine 15 (692), eade9078.

Brent, R. P. (1971). An algorithm with guaranteed convergence for finding a zero of a function. The Computer Journal 14 (4), 422–425.

Buiu, C., M. V. Putz, and S. Avram (2016). Learning the relationship between the primary structure of HIV envelope glycoproteins and neutralization activity of particular antibodies by using artificial neural networks. International Journal of Molecular Sciences 17 (10), 1710.

Conti, S. and M. Karplus (2019). Estimation of the breadth of CD4bs targeting HIV antibodies by molecular modeling and machine learning. PLoS Computational Biology 15 (4), e1006954.

Corey, L., P. B. Gilbert, M. Juraska, D. C. Montefiori, L. Morris, S. T. Karuna, S. Edupuganti, N. M. Mgodi, A. C. DeCamp, E. Rudnicki, et al. (2021). Two randomized trials of neutralizing antibodies to prevent HIV-1 acquisition. New England Journal of Medicine 384 (11), 1003–1014.

Dănăilă, V.-R. and C. Buiu (2022). Prediction of HIV sensitivity to monoclonal antibodies using aminoacid sequences and deep learning. Bioinformatics 38 (18), 4278–4285.

Follmann, D. (2018). Reliably picking the best endpoint. Statistics in Medicine 37 (29), 4374–4385.

Hake, A. and N. Pfeifer (2017). Prediction of HIV-1 sensitivity to broadly neutralizing antibodies shows a trend towards resistance over time. PLoS Computational Biology 13 (10), e1005789.

Hepler, N. L., K. Scheffler, S. Weaver, B. Murrell, D. D. Richman, D. R. Burton, P. Poignard, D. M. Smith, and S. L. Kosakovsky Pond (2014). IDEPI: rapid prediction of HIV-1 antibody epitopes and other phenotypic features from sequence data using a flexible machine learning platform. PLoS Computational Biology 10 (9), e1003842.

Huang, Y., P. B. Gilbert, R. Fu, and H. Janes (2017). Statistical methods for down-selection of treatment regimens based on multiple endpoints, with application to hiv vaccine trials. Biostatistics 18 (2), 230–243.

Julg, B. and D. Barouch (2021). Broadly neutralizing antibodies for HIV-1 prevention and therapy. In Seminars in Immunology, Volume 51, pp. 101475.

Karuna, S. T. and L. Corey (2020). Broadly neutralizing antibodies for HIV prevention. Annual Review of Medicine 71, 329–346.

LeDell, E., M. Petersen, and M. van der Laan (2015). Computationally efficient confidence intervals for cross-validated area under the ROC curve estimates. Electronic Journal of Statistics.

Magaret, C., D. Benkeser, B. Williamson, B. Borate, L. Carpp, et al. (2019). Prediction of VRC01 neutralization sensitivity by HIV-1 gp160 sequence features. PLoS Computational Biology 15 (4), e1006952.

Mahomed, S., N. Garrett, C. Baxter, Q. Abdool Karim, and S. S. Abdool Karim (2021). Clinical trials of broadly neutralizing monoclonal antibodies for human immunodeficiency virus prevention: a review. The Journal of Infectious Diseases 223 (3), 370–380.

Rawi, R., R. Mall, C.-H. Shen, S. K. Farney, A. Shiakolas, J. Zhou, H. Bensmail, T.-W. Chun, N. A. Doria-Rose, R. M. Lynch, J. R. Mascola, P. D. Kwong, and G.-Y. Chuang (2019). Accurate prediction for antibody resistance of clinical HIV-1 isolates. Scientific Reports 9 (1), 14696.

Tibshirani, R. (1996). Regression shrinkage and selection via the lasso. Journal of the Royal Statistical Society: Series B (Statistical Methodology), 267–288.

Wagh, K., T. Bhattacharya, C. Williamson, A. Robles, M. Bayne, et al. (2016). Optimal combinations of broadly neutralizing antibodies for prevention and treatment of HIV-1 clade C infection. PLoS Pathogens 12 (3), e1005520.

Wagh, K., M. Seaman, M. Zingg, T. Fitzsimons, D. Barouch, et al. (2018). Potential of conventional & bispecific broadly neutralizing antibodies for prevention of HIV-1 subtype A, C & D infections. PLoS Pathogens 14 (3), e1006860.

Walsh, S. R. and M. S. Seaman (2021). Broadly neutralizing antibodies for HIV-1 prevention. Frontiers in Immunology 12, 712122.

Williamson, B., P. Gilbert, N. Simon, and M. Carone (2021). A general framework for inference on algorithm-agnostic variable importance. Journal of the American Statistical Association (Theory & Methods).

Williamson, B. D., C. A. Magaret, P. B. Gilbert, S. Nizam, C. Simmons, and D. Benkeser (2021). Super learner prediction of nab panels (SLAPNAP): a containerized tool for predicting combination monoclonal broadly neutralizing antibody sensitivity. Bioinformatics 37 (22), 4187–4192.

Williamson, B. D., C. A. Magaret, S. Karuna, L. N. Carpp, H. C. Gelderblom, Y. Huang, D. Benkeser, and P. B. Gilbert (2023). Application of the SLAPNAP statistical learning tool to broadly neutralizing antibody HIV prevention research. iScience 26 (9).

Yoon, H., J. Macke, A. J. West, B. Foley, P. Bjorkman, et al. (2015). CATNAP: a tool to compile, analyze, and tally neutralizing antibody panels. Nucleic Acids Research 43 (W1), W213–219. doi: 10.1093/nar/gkv404.

Yu, W.-H., D. Su, J. Torabi, C. M. Fennessey, A. Shiakolas, R. Lynch, T.-W. Chun, N. Doria-Rose, G. Alter, M. S. Seaman, et al. (2019). Predicting the broadly neutralizing antibody susceptibility of the HIV reservoir. JCI Insight 4 (17).

